# Manipulation of subcortical and deep cortical activity in the primate brain using transcranial focused ultrasound stimulation

**DOI:** 10.1101/342303

**Authors:** Davide Folloni, Lennart Verhagen, Rogier B. Mars, Elsa Fouragnan, Charlotte Constans, Jean-François Aubry, Matthew F.S. Rushworth, Jérôme Sallet

**Affiliations:** Wellcome Centre for Integrative Neuroimaging (WIN), Department of Experimental Psychology, University of Oxford, Oxford, UK; Wellcome Centre for Integrative Neuroimaging (WIN), Centre for Functional MRI of the Brain (FMRIB), Nuffield Department of Clinical Neurosciences, John Radcliffe Hospital, University of Oxford; Donders Institute for Brain, Cognition and Behaviour, Radboud University Nijmegen, Nijmegen, the Netherlands; School of Psychology, University of Plymouth, UK; Institut Langevin, ESPCI Paris, PSL Research University, CNRS 7587, UMRS 979 INSERM, Université Paris Diderot, Sorbonne Paris Cité, Paris, France

## Abstract

The causal role of an area within a neural network can be determined by interfering with its activity and measuring the impact. Many current reversible manipulation techniques have limitations preventing their focal application particularly in deep areas of the primate brain. Here we demonstrate a transcranial focused ultrasound stimulation (TUS) protocol that manipulates activity even in deep brain areas: a subcortical brain structure, the amygdala (experiment 1), and a deep cortical region, anterior cingulate cortex (ACC, experiment 2), in macaques. TUS neuromodulatory effects were measured by examining relationships between activity in each area and the rest of the brain using functional magnetic resonance imaging (fMRI). In control conditions without sonication, activity in a given area is related to activity in interconnected regions but such relationships are reduced after sonication. Dissociable and focal effects on neural activity could not be explained by auditory artefacts.

## INTRODUCTION

To establish the functional role of a brain area it is necessary to examine the impact of disrupting or altering its activity. It has recently been proposed that this might be accomplished with low-intensity transcranial focused ultrasound stimulation (TUS) (Tufail et al., 2011; King et al., 2013; Yoo et al., 2011). When used over the frontal eye field in macaques, TUS leads to latency change during voluntary saccades (Deffieux et al., 2013). Comparatively little, however, is known about TUS’s impact on neural activity and if its effects persist after the stimulation has terminated. We show here that in the macaque *(Macaca mulatta)* TUS modulates neural activity and does so even in subcortical nuclei such as the amygdala and deep cortical regions such as anterior cingulate cortex (ACC). Moreover, we demonstrate a protocol that exerts an “offline” effect that lasts for an extended period of tens of minutes after an initial stimulation period of 40 s. This extended period of action is important because it means that its neural effect substantially outlasts any potential direct acoustic or somatosensory effects that might occur during the stimulation period itself (Guo et al., 2018; Sato et al., 2018). The focal impact of offline TUS in deep brain structures may underlie the specific patterns of behavioral impairment recently reported when the same protocol was used in awake behaving animals (Fouragnan et al. BioRXiv).

## RESULTS

### Stimulation of deep brain structure and resting-state fMRI recording

On each day of TUS application, a 40 s train of pulsed ultrasound (250kHz) comprising 30 ms bursts of ultrasound every 100 ms was directed to the target brain region using a single-element transducer in conjunction with a region-specific coupling cone filled with degassed water. To control for any confounds resulting from concomitant ultrasound stimulation and neural signal recording (Guo et al., 2018; Sato et al., 2018), recordings of neural activity only begun approximately 30 minutes after the end of TUS application when any potential auditory or somatosensory effects of stimulation were dissipated. We therefore refer to this stimulation protocol as an “offline” protocol.

Frameless stereotaxic neuronavigation was used to position the transducer over the target brain area taking into consideration the focal depth of the sonication (experiment 1: amygdala n=4; experiment 2: ACC n=3; relatively deep brain regions known to be interconnected and co-active during similar cognitive processes such as social cognition (Munuera et al., 2018; Noonan et al., 2014)). A single train was applied sequentially to each of the more laterally situated amygdalae, in experiment 1 and to the midline structure, ACC, in experiment 2.

The impact of TUS was determined by examining brain activity over an 80 minute period starting approximately 30 minutes after the 40 s stimulation train began (Supplementary Material). Activity was recorded not just from the stimulated site but from across the entire brain using functional MRI (fMRI). FMRI data from the stimulated animals was compared with data from an additional group of control individuals (n=9) that had received no TUS. Note that depth of anaesthesia and the delay between sedation induction and data acquisition were similar between the TUS and the control groups (0.7-0.8% and 0.7-1% range of expired isoflurane concentration, 1.53 and 2.38 hours, respectively; Supplementary Material). FMRI data were acquired at 3T under isoflurane anesthesia and preprocessed using established tools and protocols (Verhagen et al., BioRXiv; Supplementary Material). The anesthesia protocol has previously been shown to preserve regional functional connectivity measurable with fMRI (Sallet et al., 2013; Neubert et al., 2015).

Although the blood oxygen level dependent (BOLD) signal recorded with fMRI does not provide an absolute measure of activity it does provide a relative measure of activity change in relation to external events or activity recorded from other brain areas. This means that we cannot easily use BOLD to capture a measure such as activity in a brain area averaged over time. However, what we can do is to examine how BOLD responses in one area, such as the one that we are sonicating, relate to BOLD in another area using approaches similar to those employed previously (Sallet et al., 2013; Neubert et al., 2015; Vincent et al., 2007; Margulies et al., 2016; Margulies et al., 2009; Ghahremani et al., 2017; Mars et al., 2013; Shen, 2015; Shen et al., 2015; Hutchison et al., 2012).

Even at rest in the control state, BOLD activity in one area is correlated with BOLD activity in other areas and such relationships are most prominent when the areas are monosynaptically connected although some residual connectivity is mediated by indirect connections (O’Reilly et al., 2013). The pattern of activity coupling for any given area reflects its unique constellation of projections and interactions, sometimes called its “connectivity fingerprint” (Passingham et al., 2002).

### Focal effects of TUS on subcortical neural activity in the amygdala

To examine the spatial specificity of TUS effects and to investigate the capacity of TUS to stimulate subcortical structures we investigated its effects on the coupling of amygdala activity with activity in other brain areas.

If amygdala TUS affects activity in amygdala in a specific manner then what we should see is that the relationship seen at rest between the activity in amygdala and activity elsewhere will change. This does not mean that activity induced by TUS is diffused across the brain or that it is induced in one area and then “spreads” to others. Instead, to understand what impact TUS might have, we need to recall the interpretation of correlations in activity between brain areas that are found in the control state. In the control state, the relationship between activity in amygdala and activity elsewhere suggests activity in amygdala is influenced by activity in other nodes of the network the amygdala is part of and *vice versa.* If TUS dramatically modulates activity in amygdala then it may become less responsive to activity elsewhere in the brain and the relationship between activity in amygdala and elsewhere will decrease. Alternatively, if TUS strongly modulates activity in amygdala and this is associated with an increase in the activity of amygdala neurons projecting to other areas and this then results in activity change in the interconnected areas then the relationship between amygdala activity and activity elsewhere will increase. It is even possible that amygdala stimulation may lead to a pattern of change incorporating elements of both increased activity coupling with some areas and decreased coupling with other areas.

In controls, amygdala activity is coupled with activity in cingulate, ventral, and orbitofrontal cortex, striatum and the anterior temporal lobe (figs. 2a,4a). Activity coupling between the amygdala and all of these areas was reduced after amygdala TUS (non-parametric permutation test, p = 0.0020; figs. 2b,4a). These amygdala connectivity effects, however, were not found after ACC TUS, which instead left most of amygdala’s coupling pattern unaffected (non-parametric permutation test, p = 0.1346; figs.2c,4a) although not surprisingly ACC TUS led to alteration in amygdala’s coupling with ACC.

**1.**
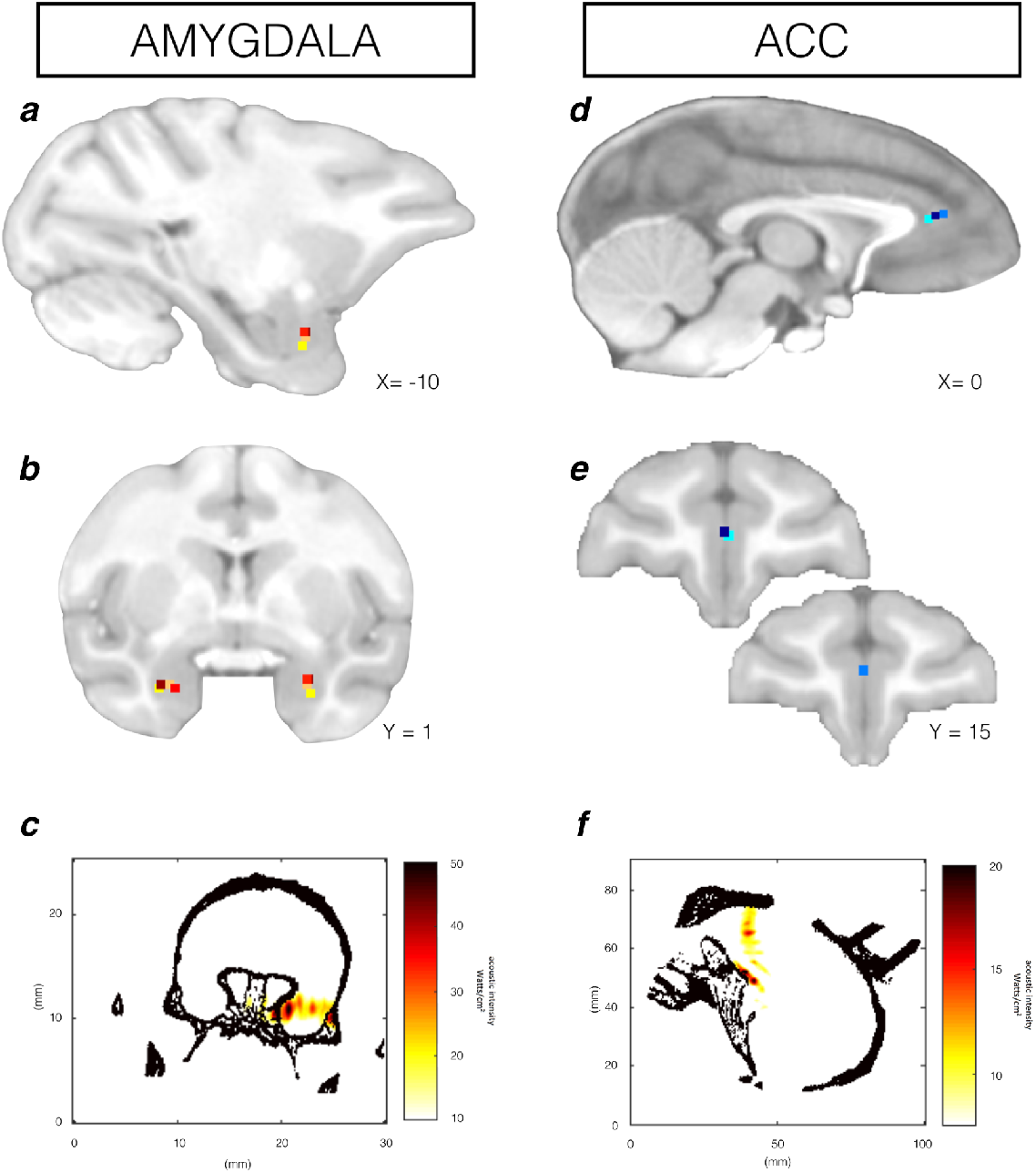
Stimulation targets. Stimulation target position is shown for each individual animal (colored dots) on sagittal and coronal views for TUS targeted at amygdala (**a-b**) and ACC (**d-e**). Acoustic intensity field (Watts/cm^2^) generated by the ultrasound beam in the brain is shown for one example animal per TUS target, amygdala (coronal plane **c**; maximum spatial peak pulse average intensity (I_sppa_) at stimulation target=51 W/cm^2^; I_spta_ = 15.3 W/cm^2^; max pressure=1.32 MPa) and ACC (sagittal plane **f**; I_sppa_ at stimulation target=17 W/cm2; I_spta_ = 5.1 W/cm2; max pressure=0.82 MPa). Note that while the target position can be delineated with accuracy in all animals in panels a, b, d, and e and activity in these areas can be examined in all subsequent figures, some slight imprecision in the simulation in panels c and f may occur when the skull model used for the simulation is applied to a given individual data set.

**2.**
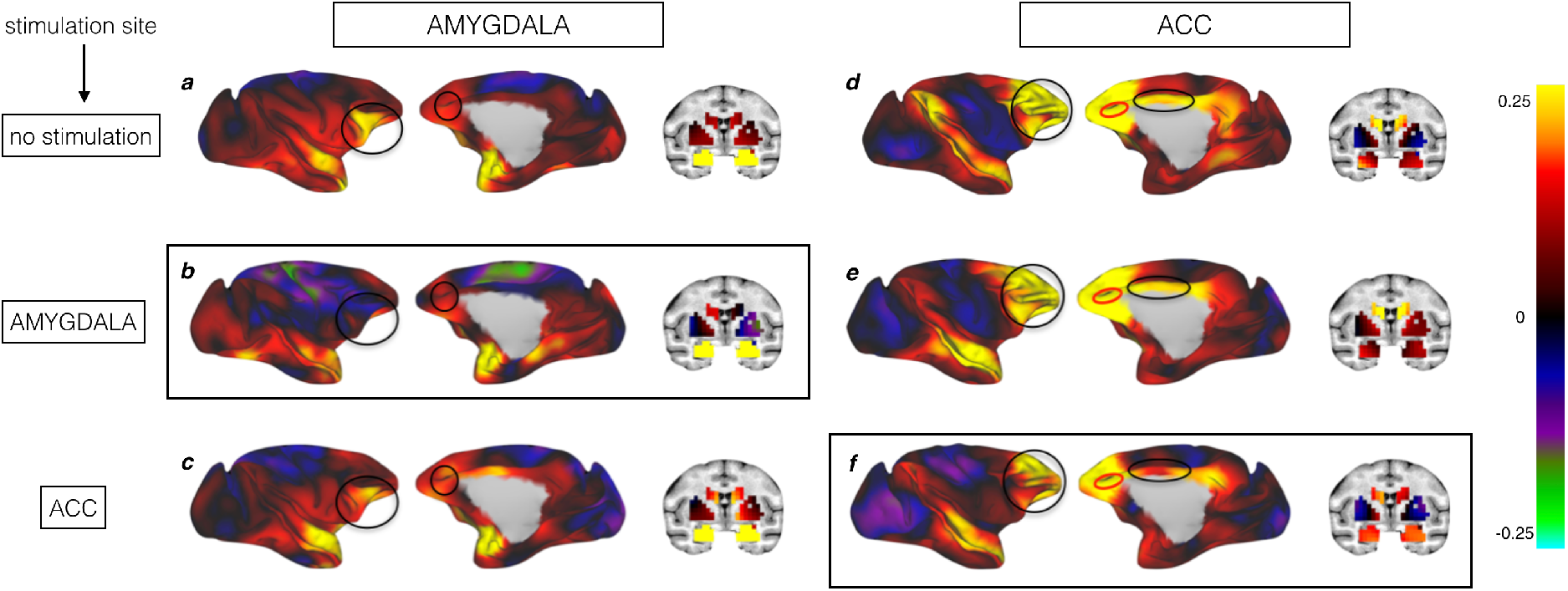
Whole-brain functional connectivity between stimulated areas and the rest of the brain. Panels a, b, and c on the left side of the figure show activity coupling between amygdala (seed masked in yellow) and the rest of the brain in no stimulation/control condition (**a**), after amygdala TUS (**b**), and after ACC TUS (**c**). Panels d, e, f show activity coupling between ACC (circled in red) and the rest of the brain in no stimulation/control condition (**d**), after amygdala TUS (**e**), and after ACC TUS (**f**). Hot colors indicate positive coupling (Fisher’s z). Functional connectivity from TUS-targeted regions are highlighted by black boxes. Each type of TUS had a selective effect on the stimulated area: amygdala coupling was strongly changed by amygdala TUS only (b) and ACC coupling was strongly changed by ACC TUS only (f). Areas showing changes in coupling with TUS-targeted regions after TUS are circled in black and compared with the other 2 control conditions.

Second, as an additional control procedure, we also investigated the coupling pattern of a control area with a very distinct constellation of projections: ventral premotor area F5c. The patterns of activity coupling between F5c and the rest of the brain that were observed both before and after amygdala TUS were virtually indistinguishable from control (fig. S1).

### Focal effects of TUS on deep cortical neural activity in anterior cingulate cortex (ACC)

To further examine the spatial specificity of TUS effects and to investigate the capacity of TUS to stimulate also deep cortical structures we investigated its effects on the coupling of ACC activity with activity in other brain areas.

In control animals at rest ACC activity was coupled with activity in strongly connected areas: dorsal, lateral, and orbital prefrontal cortex (PFC), frontal pole, mid and posterior cingulate (figs. 2d,4b). After ACC TUS the ACC coupling pattern was altered (non-parametric permutation test, p = 0.0210; fig.2f,4b). A parsimonious interpretation is that normally the activity that arises in ACC is a function of the activity in the areas that project to it, but this is no longer the case when ACC’s activity is artificially driven or diminished by TUS. Because these interactions with other areas determine the information ACC receives from elsewhere in the brain and the influence it exerts over other areas, ACC TUS should alter ACC’s computation and induce specific changes in behavior (Fouragnan et al., Bio-RXiv).

The specificity and selectivity of the effects are underscored by the results observed when 1) stimulating at another location and 2) mapping the coupling pattern of another brain area. First, after TUS over the anatomically highly connected amygdala (Amaral and Price, 1984), there was minimal change in ACC coupling with other brain areas, beyond its coupling with the amygdala itself and caudal orbitofrontal areas. This suggests ACC-caudal orbitofrontal coupling is mediated by amygdala (figs. 2e,4b). However, because the caudal orbitofrontal cortex effect was so circumscribed this effect of amygdala TUS did not lead to a significant change in ACC connectivity in general (non-parametric permutation test, p = 0.0744). Importantly, there was a significant difference between ACC and amygdala TUS effects (non-parametric permutation test, p = 0.0428).

Just as amygdala TUS did not affect the activity relationships between F5c and other brain regions, ACC TUS also did not affect F5c’s coupling (fig.S1).

### Focality of TUS

Our major focus in the current investigation is primarily on the possibility of altering activity in deep brain structures with ultrasound and the sonication parameters adopted here have been optimized with this aim in mind. In future experiments it will be possible to manipulate the ultrasound’s features to enhance the spatial focality of any effects that we find. Nevertheless, it is obviously of interest to examine briefly the focality that is obtained with the current sonication parameters. An additional set of analyses was, therefore, also conducted. Instead of examining the pattern of activity coupling between the stimulated areas and the rest of the brain in a control state and after TUS, these analyses focused on the areas surrounding the stimulated areas or located between the stimulation cone and the target area (figs.3, S2).

**3.**
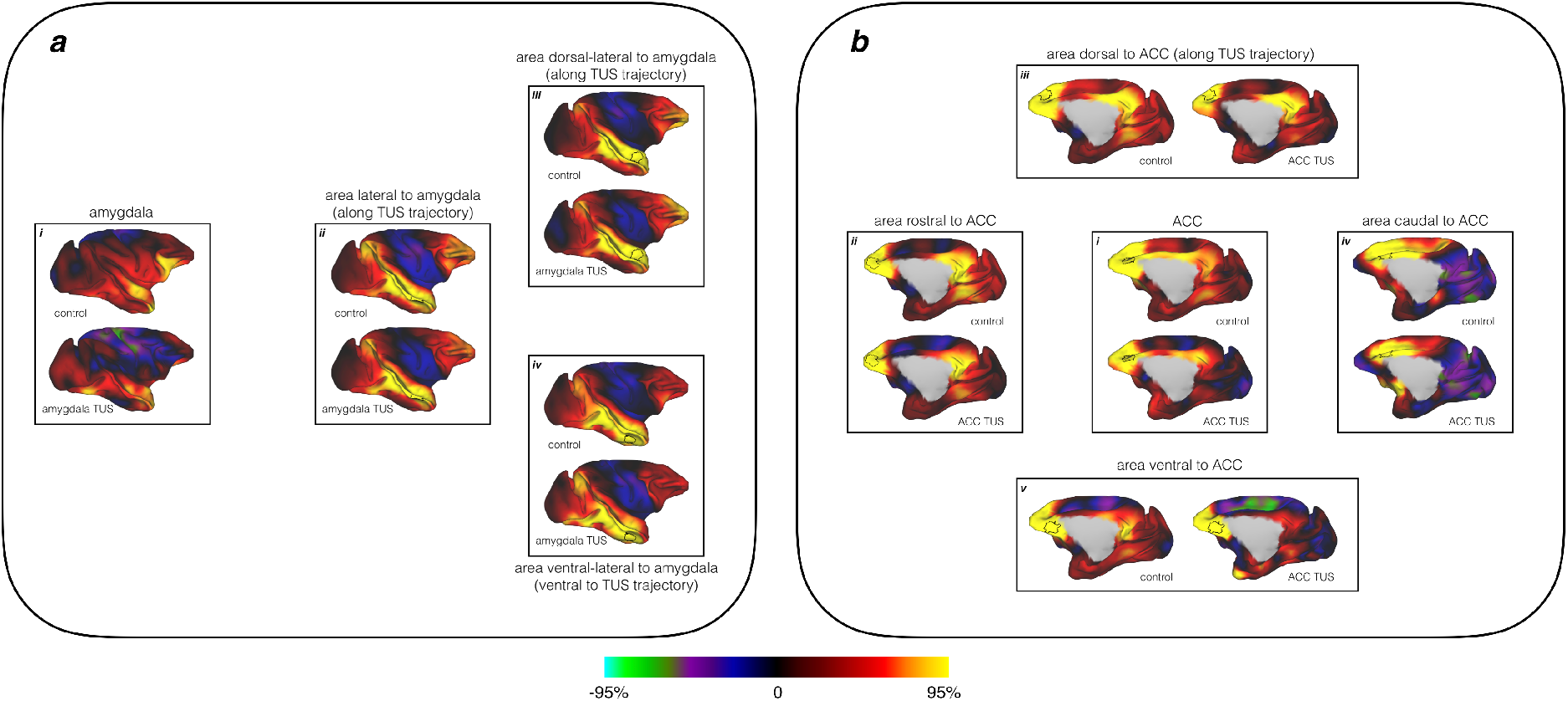
Whole-brain activity coupling of the areas neighboring the stimulated region in the control condition and after amygdala and ACC TUS. *Panel a;* as also shown in figure 2, the whole brain coupling of the amygdala target region (sub-panel *i;* the outline in black in all cases indicates the regions for which the whole-brain connectivity is shown) is significantly different in the control condition and when TUS is applied to amygdala. Hot colors indicate positive coupling (Fisher’s z). Sub-panels *ii, iii, iv* show the activity coupling of regions along *(ii,iii)* or immediately surrounding the trajectory of the ultrasound stimulation beam (*iv*), in the control condition and after amygdala TUS. There were no changes in the coupling of these regions and the rest of the brain. This is true for the regions through which the stimulation trajectory passed in the fundus of the superior temporal sulcus *(ii)* and the superior temporal gyrus *(iii)* or in the immediately adjacent inferior temporal gyrus (*iv*). *Panel b;* the whole brain coupling of the ACC target region (*i*) is significantly different in the control condition and when TUS is applied to ACC. Sub-panels ii, *iii, iv* and *v* show the activity coupling of regions near the ACC target in both the ACC TUS and control conditions. This includes areas located along the trajectory of the ultrasound stimulation beam such as (*iii*) the area in between the transducer and the target region in ACC and the area on the other side of the target region (*v*) as well as areas immediately rostral (*ii*) and caudal (*iv*) to the target region. Some changes in coupling can be seen along the stimulation trajectory in the area just ventral to the target (*v*) and also in an area which is unlikely to have been hit directly by the ultrasound beam (*iv*). These areas are strongly anatomically connected with the targeted area.

**4.**
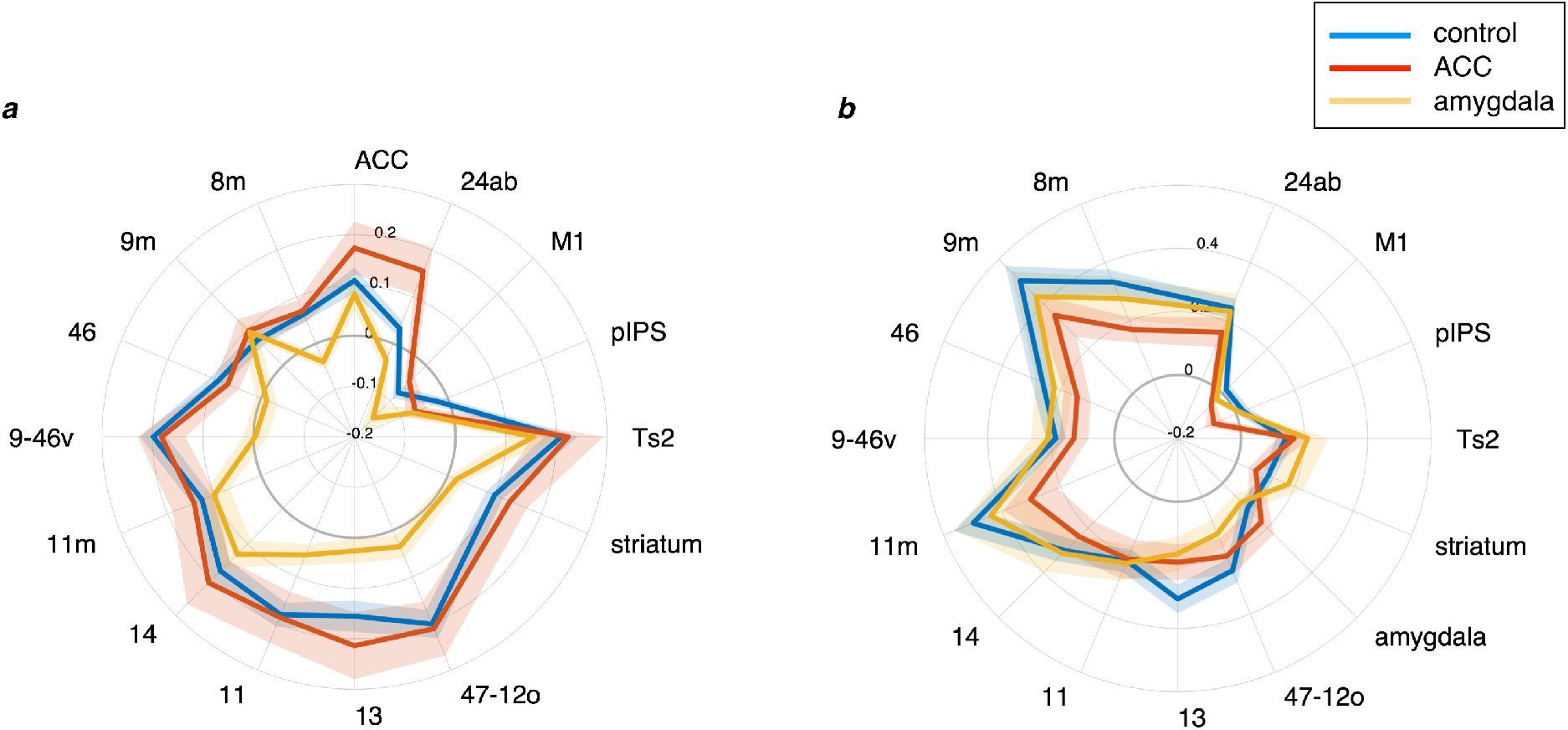
ACC and amygdala connectivity fingerprints after stimulation. The lines in the left panel indicates the strength of activity coupling between amygdala (**a**) and other brain areas labelled on the circumference in control animals (blue), after amygdala TUS (yellow), and after ACC TUS (red). The lines in the right panel show activity coupling between ACC (**b**) and the rest of the brain in control animals (blue), after ACC TUS (red), and after amygdala TUS (yellow). Each type of TUS had a selective effect on the stimulated area: amygdala coupling was strongly affected by amygdala TUS (the yellow line is closer to the center of the panel than the blue line) and ACC coupling was strongly disrupted by ACC TUS (the red line is closer to the center of the panel than the blue line). Standard error of the mean is indicated by shading around each line.

In the first set of analyses we measured activity coupling before and after amygdala TUS in three control areas located along the trajectory of the ultrasound beam (fig.3a, sub-panels *ii* and *iii*) or just ventral to it (fig.3a, sub-panel *iv*). Interestingly, when TUS was directed at the amygdala, there were no major changes in the activity coupling of areas situated on the trajectory of the ultrasound beam such as the superior temporal gyrus (fig.3a, sub-panel *iii*) and fundus of the superior temporal sulcus (fig.3a, sub-panel *ii*) or of the inferior temporal gyrus which was just ventral to the TUS trajectory (fig.3a, sub-panel *iv*). Similarly, we measured changes in coupling between four control areas and the rest of the brain in a control state and after TUS targeted to ACC. These areas were located in between the transducer and the ACC target (fig.3b, sub-panel iii), on the other side of the target region (fig.3b, sub-panel v) as well as areas immediately rostral (fig.3b, sub-panel ii) and caudal (fig.3b, sub-panel iv) to the target region. ACC sonication had little effect on the region between the target and the transducer (fig.3biii) and a region just anterior to the target (fig.3bii) suggesting, once again, a considerable degree of focality in the effect on neural activity. Unlike what we had observed for the amygdala, however, we noticed that there were some changes in coupling between some of the control regions surrounding ACC and the rest of the brain. These effects can be seen in one of the areas most at risk of being stimulated because it lay along the stimulation trajectory just ventral to the target (fig.3b, v), in a region where the acoustic waves are likely to be reflected on the orbital bone (fig.1f). However, the same was also true of an area just posterior (fig.3b, iv) to the target region (fig.3b, *ii*), despite the fact that it was unlikely to have been directly sonicated. An additional analysis was conducted to test whether changes in adjacent areas were due to their spatial proximity to the target area or the anatomical connections they shared with the target area. When we investigated an area - the SMA - which is at a similar Euclidean distance from the target ACC region (fig.S2a) as the more caudal cingulate area 24ab, but less strongly connected to it, there were no changes (fig.S2b) in the way in which its activity was coupled with that in other brain areas. This was in contrast with changes in activity coupling of area 24ab (fig.S2c), which is more strongly connected to the ACC target compared to SMA (fig.S2b).

One possible explanation for the difference between our two stimulation sites might be related to the nature of the connections between the control regions and the sonicated region. While the different areas of the medial prefrontal cortex are strongly interconnected (Yeterian et al., 2012), the amygdala connections with the control regions were limited to the most lateral and dorsal nuclei (Amaral et al., 1992). Another potential explanation might be related to the difference in the transmission of the ultrasound waves between the two sites. For instance it is noticeable that while ultrasound intensity was greatest at the target ACC area (fig.1f) some changes in activity coupling after ACC TUS were also present in the more ventral region (fig.3b, v). This later issue may be overcome in the future by targeting the sonication site more precisely using higher frequency ultrasound (500kHz) or multiple beams on different trajectories that intersect at the target location.

### Effect of amygdala and ACC TUS on the auditory system

It has recently been suggested that TUS’s impact on neural activity is mediated by its auditory impact (Guo et al., 2018; Sato et al., 2018). Several considerations suggest that it might not be possible to explain away the current findings as the result of an auditory artefact. First, the auditory impact of TUS is likely a function of specific features of its frequency and pulse type. Second, the auditory stimulation associated with the TUS application ceased after the 40 s sonication period but the neural activity measurements were initiated tens of minutes later. Third, TUS of each area, ACC and amygdala, had specific effects that were distinct to one another. The only amygdala activity relationship affected by ACC TUS was that between amygdala and ACC and the only ACC activity relationship affected by amygdala TUS was that between ACC and amygdala.

Nevertheless, we also carried out a fourth line of inquiry and examined whether it is plausible that an auditory effect could have mediated the TUS effects on amygdala and ACC. To quantify this probability, we correlated any TUS effects on primary auditory cortex (A1) connectivity with TUS effects on the targeted regions (fig.S3a). TUS effects on the auditory cortex after both amygdala (r=0.1084, p=0.7007) and ACC (r=0.1474, p=0.6000) sonication are unrelated to the TUS effects at each target site and are therefore unlikely to have mediated effects seen at the stimulation sites. However, it is possible that TUS over amygdala or ACC had an impact on A1 connectivity separately from its impact on the stimulated sites themselves (fig.S3b). While A1 connectivity is not impacted by ACC TUS (fig.S3a, non-parametric permutation test, p=0.6871), amygdala TUS did have a significant impact on A1 connectivity (fig.S3a, non-parametric permutation test, p = 0.0002). Closer inspection revealed that this was due to a diminution solely in A1’s interactions with the amygdala itself and two areas with which the amygdala is itself strongly connected with: ACC and orbitofrontal cortex. Differential effects of ACC and amygdala TUS on A1 connectivity might be driven by some direct, albeit weak, connections of amygdala with A1 (Yukie, 2002). Similarly, given amygdala’s strong connections to ACC and orbitofrontal cortex, it is perhaps not surprising that amygdala sonication might affect A1’s interactions with them. Importantly, these circumscribed effects on A1 connectivity are not predictive of the effects elsewhere. In summary, the alteration seen in the A1 fingerprint is a poor match to the alteration seen in the amygdala fingerprint after amygdala TUS or in the ACC fingerprint after ACC TUS.

## DISCUSSION

In these investigations we combined TUS with resting-state fMRI to examine the impact of modulating activity in subcortical and deep cortical areas of the primate brain. Experiments 1 and 2 revealed dissociable effects of amygdala and ACC TUS. The dissociable nature of the effects and the fact that they were observed more than an hour after the 40 s stimulation period suggests they are not mediated by the stimulation’s auditory impact (Guo et al., 2018; Sato et al., 2018). In each case effects were apparent as reductions in activity coupling between the stimulated area and other regions with which it is normally interconnected; after TUS, a brain area’s activity appears to be driven less by activity in the areas with which it is connected and more so by the artificial modulation induced by TUS.

The impact that TUS exerts on the auditory system is likely to depend on the precise details of the sonication frequency and pulse type and might be specific to other features of the preparation such as anesthesia level (Airan and Pauly, 2018). In addition to the offline protocol used here, it may be possible to develop other procedures to diminish TUS’s auditory impact. It is also possible that TUS may act not simply by immediately inducing or reducing activity in neurons but by modulating their responsiveness to other neural inputs.

Several molecular mechanisms describing how low-intensity ultrasound stimulation modulates neuronal activity have been suggested. However, recent investigations on the interactions between sound pressure waves and brain tissue suggest that ultrasound primarily exerts its modulatory effects through a mechanical action on cell membranes, notably affecting ion channel gating (Kubanek et al., 2016; Prieto et al., 2013; Tyler et al., 2008). While the precise mechanism is being determined (Kubanek, 2018; Kubanek et al., 2018; Tyler et al., 2018) the current results suggest TUS may be suitable as a tool for focal manipulation of activity in many brain areas in primates. Specifically, they show that TUS may even be used to manipulate activity in subcortical structures in monkeys.

TUS’s capacity to stimulate subcortical and deep cortical areas in primates, therefore, opens the prospect of advanced non-invasive causal brain mapping. To date, non-invasive manipulation of brain activity in humans can be done reversibly only using neuromodulation methods such as transcranial magnetic stimulation and transcranial current stimulation. However, the spatial resolution of some of these techniques is limited (Walsh and Cowey, 2000; Dayan et al., 2013). Even more critically, application of these techniques is constrained to the surface of the brain as their efficacy falls off rapidly with depth.

In another recent study we have shown that TUS of the type used here causes no permanent damage to tissue on histological analysis (Verhagen et al., bioRxiv). While such results are encouraging, it will only be possible to extend the technique to humans if care is taken with the assessment of each new protocol that is devised. For example, it is notable that the neural effects of the stimulation protocol used here are sustained over a period of time that is substantially longer than that used in many laboratory-based cognitive neuroscience experiments (Verhagen et al., bioRxiv). In addition, the need for care is also underlined by the impact of sonication on meningeal compartment (Verhagen et al., bioRxiv). This will probably mean that the impact of TUS to a brain area is best assessed by comparison to the impact of TUS to a control site.

In summary, based on the results reported here, TUS can be used to transiently and reversibly alter neural activity in subcortical and deep cortical areas with high spatial specificity. To date, it is the most promising neuromodulatory technique to reach areas deep below the dorsolateral surface of the brain in a non-invasive and focal manner, thereby providing it with the potential for causally mapping brain functions within and across species. While it may currently lack the capacity to target specific neurons, as do some optogenetic and chemogenetic techniques (Khatoun et al., 2017; Sternson and Roth, 2014; Tang et al., 2018; Yizhar et al., 2011), it may provide a method for investigating brain areas that may make it suitable for use with primate species, which are rarely investigated with such techniques even though many brain areas are particularly well developed or only present in primates (Passingham and Wise, 2012). With care it may even be possible to employ offline TUS protocols in investigations of human brain function.

## Acknowledgements

Funding for this study was provided by a Wellcome Senior Investigator Award (WT100973AIA), MRC programme grant (MR/P024955/1), MRC programme grant (G0902373), Wellcome Trust Henry Dale Fellowship (105651/Z/14/Z), Wellcome Trust UK Grant (105238/Z/14/Z), The Wellcome Centre for Integrative Neuroimaging (203139/Z/16/Z), BBRSC grant (BB/N019814/1), NWO grant (452-13-015), Bettencourt Schueller Foundation and the “Agence Nationale de la Recherche” under the program ‘‘Future Investments’’ (ANR-10-EQPX-15).

## Contributions

D.F. and J.S. designed the experiment; D.F. and J.S. collected the data; D.F., L.V., M.F.S.R., and J.S. analyzed the data; L.V. and R.B.M. contributed analysis tools; J.F.A., C.C, and D.F. contributed to the ultrasound modelling; D.F., L.V., E.F., M.F.S.R., J.F.A., and J.S. wrote the manuscript.

## Declaration of Interests

The authors have no competing interests to declare.

## METHOD DETAILS

### Ultrasound stimulation

A single-element ultrasound transducer (H115-MR, diameter 64 mm, Sonic Concept, Bothell, WA, USA) with a 51.74 mm focal depth was used with region-specific coupling cones filled with degassed water and sealed with a latex membrane (Durex) to assess TUS of ACC (experiment 1; n=3) and amygdala (experiment 2; n=4) (fig.1). The ultrasound wave frequency was set to the 250 kHz resonance frequency and 30 ms bursts of ultrasound were generated every 100 ms (duty cycle 30%) with a digital function generator (Handyscope HS5, TiePie engineering, Sneek, the Netherlands). Overall, the stimulation lasted for 40 s. A 75-Watt amplifier (75A250A, Amplifier Research, Souderton, PA) was used to deliver the required power to the transducer. A TiePie probe (Handyscope HS5, TiePie engineering, Sneek, The Netherlands) connected to an oscilloscope was used to monitor the voltage delivered. The recorded peak-to-peak voltage was constantly maintained throughout the stimulation. Voltage values per session ranged from 128 to 134V. It corresponded to a peak negative pressure ranging from 1.15 to 1.27MPa respectively as measured in water with an in house heterodyne interferometer (Constans et al., 2017). The acoustic wave propagation of our focused ultrasound protocol was simulated at 130 V peak-to-peak voltage using finite element models of an entire monkey head to obtain estimates for the pressure amplitude, peak intensity, and spatial distribution (Constans et al., 2017). 3D maps of the skull were extracted from a monkey CT scan (0.36 mm isotropic resolution). Based on these numerical simulations, the maximum spatial peak pulse average intensity (Isspa) at the acoustic focus target was estimated to be 51 W/cm^2^ (spatial peak temporal average intensity (I_spta_)=15.3 W/cm2) in the amygdala and 17 W/cm2 (I_spta_=5.1 W/cm2) in ACC with a maximum pressure of 1.32 MPa in amygdala and 0.82 MPa in ACC. One train was applied to each of the more laterally situated amygdalae but a single train was applied to the midline structure (ACC) in experiments 1 and 2 respectively.

Each individual animal’s structural magnetic resonance (MRI) image was registered to its head with a frameless stereotaxic neuronavigation system (Rogue Research, Montreal, CA). By recording the positions of both the ultrasound transducer and the head with an infrared tracker it was then possible to co-register the ultrasound transducer with respect to the MRI scan of the brain to position the transducer over the targeted brain region, either ACC (Procyk et al., 2016) (MNI coordinates x = 0, y = 15, z = 6) or amygdala (MNI coordinates x = −10, y = 1, z = −11; x = 9, y = 1, z = −11). The ultrasound transducer / coupling cone montage was placed directly onto previously shaved skin on which conductive gel (SignaGel Electrode; Parker Laboratories Inc.) had been applied to ensure ultrasonic coupling between the transducer and the animal’s head. In the non-stimulation condition (control), all procedures (anaesthesia, pre-scan preparation, fMRI scan acquisition and timing), with the exception of actual TUS, mirrored the TUS sessions.

### Macaque rs-fMRI Data Acquisition

Resting state fMRI (rs-fMRI) and anatomical MRI scans were collected for 11 healthy macaques *(Macaca mulatta)* (two females; rs-fMRI from nine animals were acquired under no stimulation; rs-fMRI from three animals were acquired post ACC TUS; rs-fMRI from four animals were acquired post amygdala TUS; age: 7.3 years, weight: 10.3 kg) under inhalational isoflurane anesthesia using a protocol which was previously proven successful (Noonan et al., 2014; Neubert et al., 2015) in preserving whole-brain functional connectivity as measured with BOLD signal. In the case of the TUS conditions, fMRI data collection began only after completion of the TUS train (delay between ultrasound stimulation offset and scanning onset: 37.5 minutes; SEM: 2.21 minutes). Anesthesia was induced using intramuscular injection of ketamine (10 mg/kg), xylazine (0.125 – 0.25 mg/kg), and midazolam (0.1 mg/kg). Macaques also received injections of atropine (0.05 mg/kg, intramuscularly), meloxicam (0.2 mg/kg, intravenously), and ranitidine (0.05 mg/kg, intravenously). To block peripheral nerve stimulation, 15 minutes before placing the macaque in the stereotaxic frame local anaesthetic (5% lidocaine/prilocaine cream and 2.5% bupivacaine) was also administered via subcutaneous injection around the ears. The anesthetized animals were placed in an MRI-compatible stereotactic frame (Crist Instruments) in a sphinx position and placed in a horizontal 3T MRI scanner with a full-size bore. Scanning commenced 1.53 hours (SEM: 4 minutes) and 2.38 hours (SEM: 4 minutes) after anesthesia induction in TUS and control sessions, respectively. In both cases data collection commenced when the clinical peak of ketamine had passed. Anesthesia was maintained, in accordance with veterinary recommendation, using the lowest possible concentration of isoflurane to ensure that macaques were anesthetized. The depth of anesthesia was assessed and monitored using physiological parameters (heart rate and blood pressure, as well as clinical checks before the scan for muscle relaxation). During the acquisition of the functional data, the inspired isoflurane concentration was in the range 0.8 – 1.1%, and the expired isoflurane concentration was in the range 0.7-1%. Isoflurane was selected for the scans as it was previously demonstrated to preserve rs-fMRI networks (Neubert et al., 2015; Mars et al., 2013; Vincent et al., 2007). Macaques were maintained with intermittent positive pressure ventilation to ensure a constant respiration rate during the functional scan, and respiration rate, inspired and expired CO2, and inspired and expired isoflurane concentration were monitored and recorded using VitalMonitor software (Vetronic Services Ltd.). Core temperature and SpO2 were also constantly monitored throughout the scan.

A four-channel phased-array coil was used for data acquisition (Dr. H. Kolster, Windmiller Kolster Scientific, Fresno, CA, USA). Whole-brain BOLD fMRI data were collected from each animal for up to 78 minutes. All fMRI data were collected using the following parameters: 36 axial slices; inplane resolution, 2 × 2 mm; slice thickness, 2 mm; no slice gap; TR, 2000 ms; TE, 19 ms; 800 volumes per run. A minimum period of 10 days elapsed between sessions.

A structural scan (average over up to three T1-weighted structural MRI images) was acquired for each macaque in the same session as the functional scans, using a T1-weighted magnetization-prepared rapid-acquisition gradient echo sequence (0.5 × 0.5 × 0.5 mm voxel resolution).

All recording and stimulation procedures were conducted under licenses from the United Kingdom (UK) Home Office in accordance with The Animals (Scientific Procedures) Act 1986 and with the European Union guidelines (EU Directive 2010/63/EU).

### Macaque rs-fmri data preprocessing, and analysis

The preprocessing and analysis of the MRI data was designed to follow the HCP Minimal Processing Pipeline (Glasser et al., 2013), using tools of FSL (https://fsl.fmrib.ox.ac.uk/fsl/fslwiki), HCP Workbench (https://www.humanconnectome.org/software/connectome-workbench), and the Magnetic Resonance Comparative Anatomy Toolbox (MrCat; www.neuroecologylab.org). The processing pipeline has been validated and described in full (Verhagen et al., BioRxiv).

The T1w images were processed in an iterative fashion cycling through brain-extraction (BET) (Smith, 2002), RF bias-field correction, and linear and non-linear registration (FLIRT and FNIRT) (Jenkinson and Smith, 2001; Jenkinson et al., 2002) to the macaca mulatta F99 atlas(Van Essen, 2002; Van Essen and Dierker, 2007). The application of robust and macaque-optimised versions of BET and FAST (Zhang et al., 2001) also provided segmentation into grey matter, white matter, and cerebral spinal fluid compartments. Segmentation of subcortical structures was obtained by registration to the D99 atlas (Reveley et al., 2017).

The first 5 volumes of the functional EPI datasets were discarded to ensure a steady RF excitation state. EPI timeseries were motion corrected using MCFLIRT. Given that the animals were anesthetized and their heads were held in a steady position, any apparent image motion, if present at all, is caused by changes to the B0 field, rather than by head motion. Accordingly, the parameter estimates from MCFLIRT can be considered to be ‘B0-confound parameters’ instead. Each timeseries was checked rigorously for spikes and other artefacts, both visually and using automated algorithms; where applicable slices with spikes were linearly interpolated based on temporally neighboring slices. Brain extraction, bias-correction, and registration was achieved for the functional EPI datasets in an iterative manner, similar to the preprocessing of the structural images with the only difference that the mean of each functional dataset was registered to its corresponding T1w image using rigid-body boundary-based registration (FLIRT). EPI signal noise was reduced both in the frequency and temporal domain. First, the functional time series were high-pass filtered at 2000s. Temporally cyclical noise, for example originating from the respiratory apparatus, was removed using band-stop filters set dynamically to noise peaks in the frequency domain. Remaining temporal noise was described by the mean time course and first two subsequent principal components of the white matter (WM) and cerebral spinal fluid (CSF) compartment (considering only voxels with a high posterior probability of belonging to the WM or CSF, obtained in the T1w image using FAST). The B0-confound parameter estimates were expanded as a second degree Volterra series to capture both linear and non-linear B0 effects. Together the WM and CSF expanded B0 confound parameters were regressed out of the BOLD signal for each voxel.

The cleaned time course was then low-pass filtered with a cut-off at 10 seconds. The cleaned and filtered signal was projected from the conventional volumetric representation (2mm voxels) to the F99 cortical surface (~1.4mm spaced vertices) using Workbench command “myelin-style” mapping, while maintaining the subcortical volumetric structures. The data was spatially smoothed using a 3mm FWHM gaussian kernel, while taking into account the folding of the cortex and the anatomical boundaries of the subcortical structures. Lastly, the data were demeaned to prepare for functional connectivity analyses.

To represent subject effects, the timeseries from the three runs were concatenated to create a single timeseries per animal per intervention (control, ACC TUS, amygdala TUS). To represent group effects the run-concatenated timeseries of all animals were combined using a group-PCA approach (Smith et al., 2014) that was set to reduce the dimensionality of the data.

To construct a region-of-interest (ROI) for ACC, a circle of 4mm radius was drawn on the cortical surface around the point closest to the average stimulation coordinate (fig.1), in both the left and the right hemisphere. The same procedure was used to define other bilateral cortical regions of interest, based on literature coordinates (Neubert et al., 2015; Sallet et al., 2013; Neubert et al., 2014), to serve as targets for the fingerprint analyses (fig.3).

Coupling between the activity of each region of interest and the rest of the brain was estimated by calculating the Fisher’s z-transformed correlation coefficient between each point in the ROI and all other datapoints. The resulting “connectivity-maps” were averaged across all points in the ROI, across both hemispheres. Accordingly, the final maps represent the average coupling of a bilateral ROI with the rest of the brain. The fingerprints are obtained by extracting the average coupling with each target ROI and averaging across the two hemispheres. Statistical inference on the fingerprints was performed by using non-parametric permutation tests on cosine similarity metrics describing how similar or dissimilar pairs of fingerprints are(Mars et al., 2016; Verhagen et al., BioRxiv). In contrast to conventional parametric tests, this approach does not rely on assumptions about the shape of the distribution but will acknowledge dependencies between target ROIs in the fingerprint; as such this approach will avoid inflation of type I error. For each test we ran 10,000 permutations to accurately approximate the true probability of rejecting the null-hypothesis of permutable conditions.

## SUPPLEMENTAL INFORMATION

**Supplementary figure 1.**
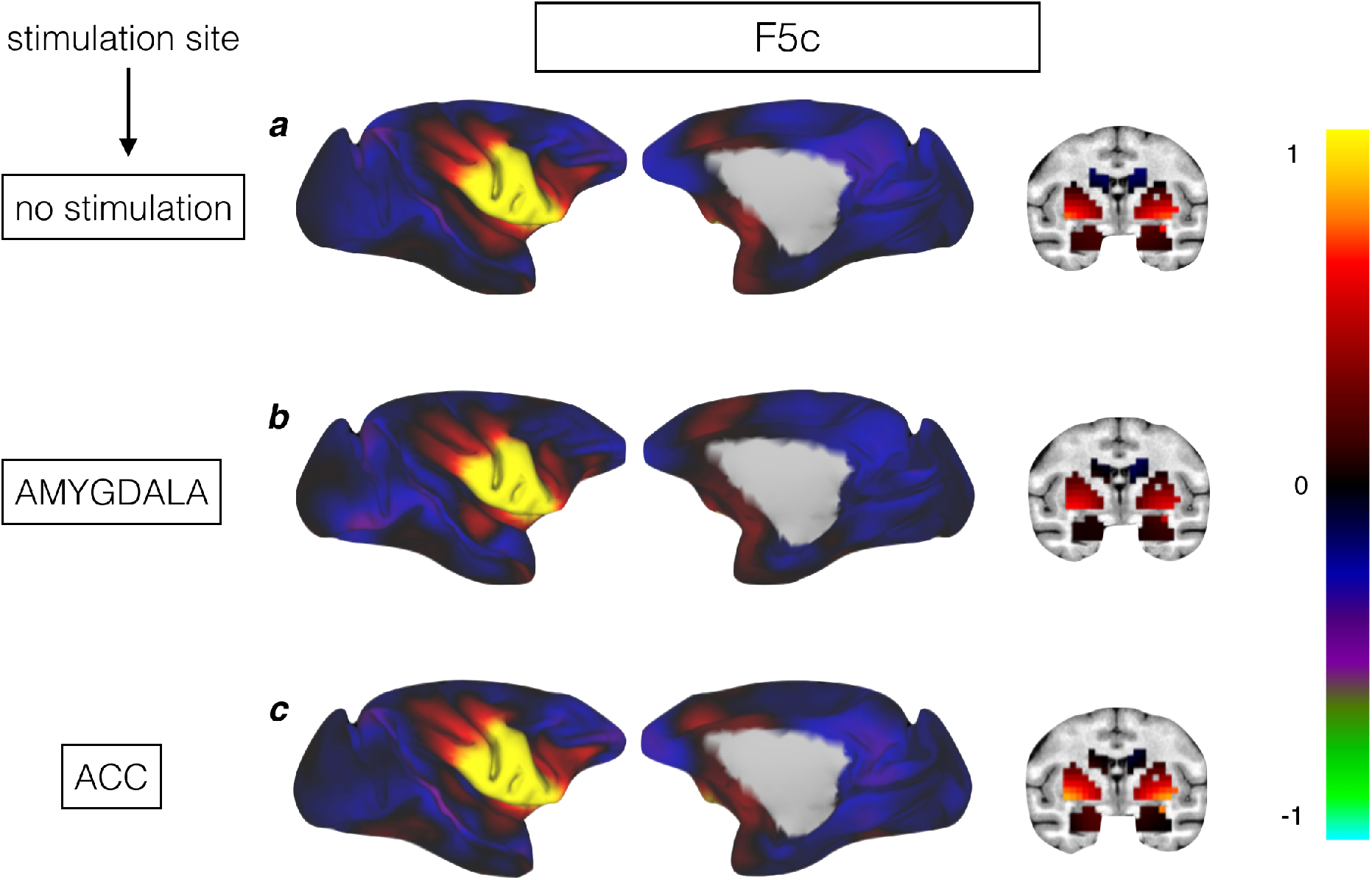
Whole-brain functional connectivity between stimulated and not stimulated areas with the rest of the brain. Panels a, b, and c show activity coupling between a control area, the caudal ventral premotor area F5c, and the rest of the brain in no stimulation/control condition (**a**), after amygdala TUS (**b**), and after ACC TUS (**c**). Hot colors indicate positive coupling (Fisher’s z). Compared to a no stimulation condition (a), neither amygdala TUS nor ACC TUS (b,c) affected the whole-brain coupling activity of F5c which has weak anatomical connections with ACC and amygdala.

**Supplementary figure 2.**
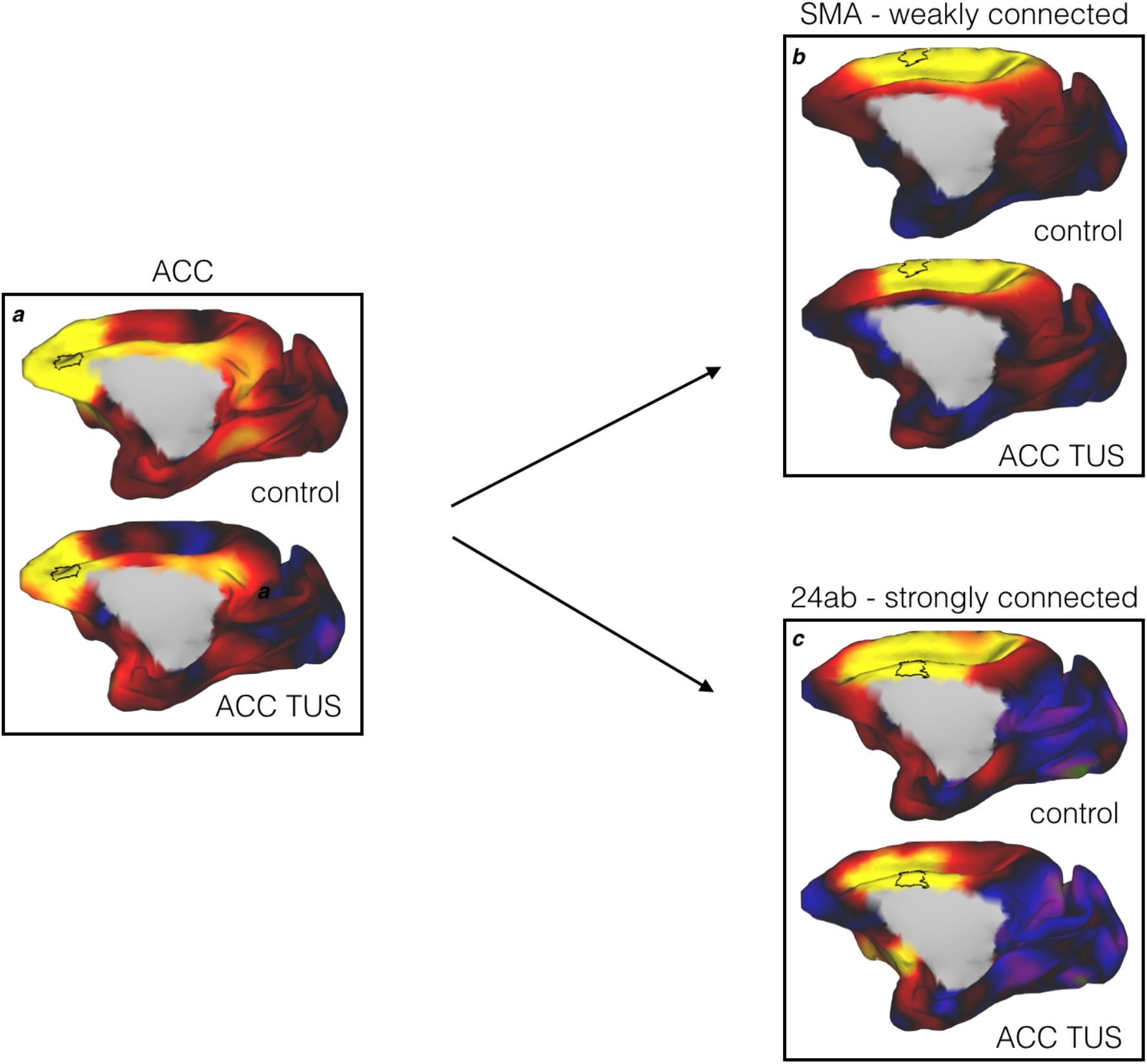
Effects of TUS on regions outside the target area are mediated by the strength of anatomical connectivity rather than a result of spatial proximity. Panels a, b, and c of the figure show whole-brain activity coupling in control and ACC TUS conditions for ACC (a) and two areas at an equal Euclidian distance from the ACC target region: SMA (b) and area 24ab (c). Hot colors indicate positive coupling (Fisher’s z). The activity coupling of SMA, an area weakly connected with the target area ACC, and the rest of the brain is predominantly preserved after TUS. However, the whole-brain coupling of an area more strongly connected with the target ACC region (Hoesen et al., 1993), area 24b, is more influenced by TUS to the ACC target region.

**Supplementary figure 3.**
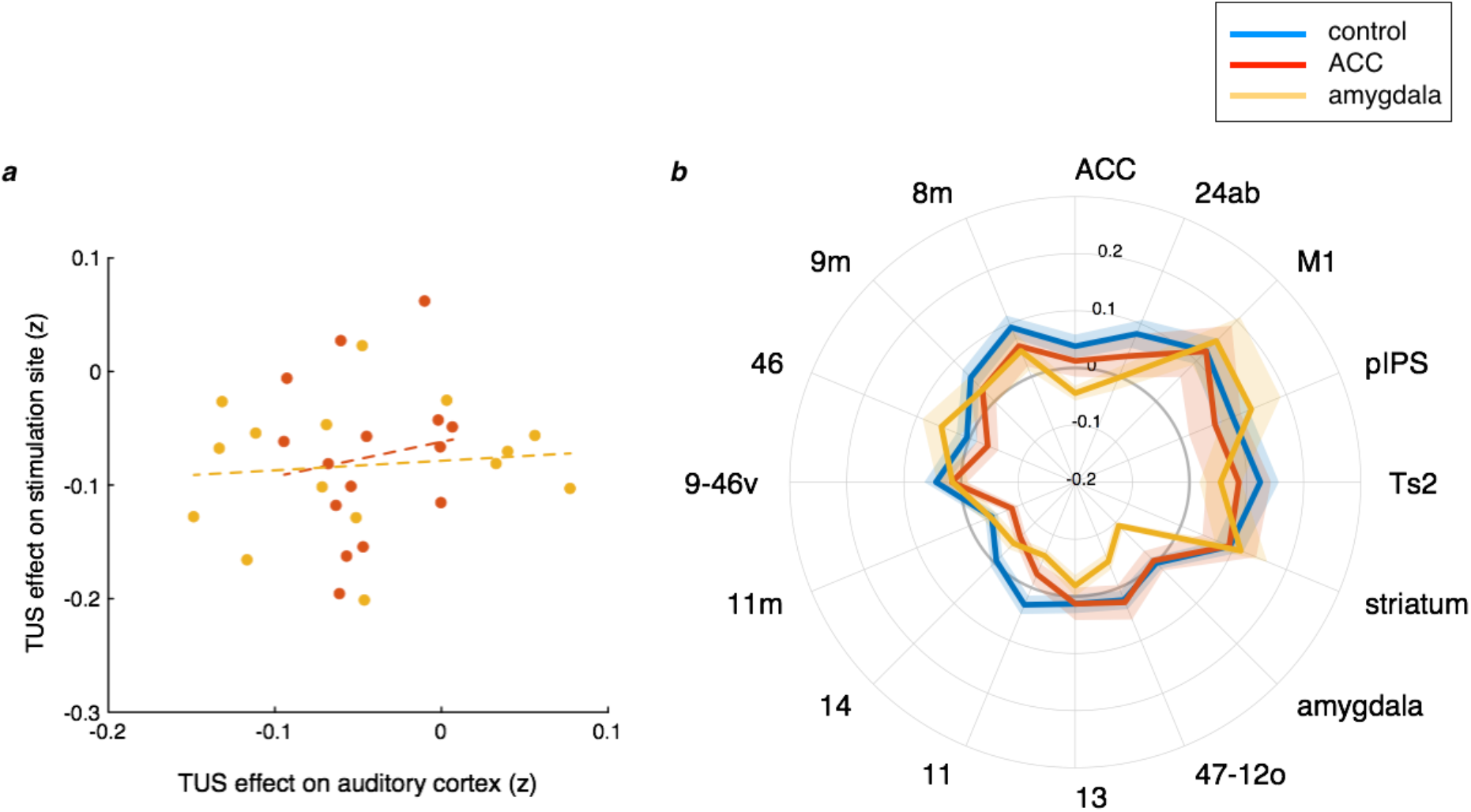
Effect of amygdala and ACC TUS on the functional coupling of primary auditory cortex. *Panel a;* ACC TUS (red line) had no effects on the functional coupling of A1. Amygdala TUS (yellow line) affected the relationship between A1’s activity and activity in several areas that are linked to the A1 via the amygdala including the amygdala itself, lateral orbitofrontal cortex area 47/12o and ACC. *Panel b;* TUS effects on the auditory cortex after neither amygdala (yellow) nor ACC (red) cannot explain the TUS effect on the respective stimulation sites.

